# The Dynamic Influence of Linker Histone Saturation within the Poly-Nucleosome Array

**DOI:** 10.1101/2020.09.20.305581

**Authors:** Dustin C. Woods, Francisco Rodríguez-Ropero, Jeff Wereszczynski

## Abstract

Linker histones bind to nucleosomes and modify chromatin structure and dynamics as a means of epigenetic regulation. Biophysical studies have shown that chromatin fibers can adopt a plethora of conformations with varying levels of compaction. Linker histone condensation, and its specific binding disposition, has been associated with directly tuning this ensemble of states. However, the atomistic dynamics and quantification of this mechanism remains poorly understood. Here, we present molecular dynamics simulations of octa-nucleosome arrays, based on a cryo-EM structure of the 30-nm chromatin fiber, with and without the globular domains of the H1 linker histone to determine how they influence fiber structures and dynamics. Results show that when bound, linker histones inhibit DNA flexibility and stabilize repeating tetra-nucleosomal units, giving rise to increased chromatin compaction. Furthermore, upon the removal of H1, there is a significant destabilization of this compact structure as the fiber adopts less strained and untwisted states. Interestingly, linker DNA sampling in the octa-nucleosome is exaggerated compared to its mono-nucleosome counterparts, suggesting that chromatin architecture plays a significant role in DNA strain even in the absence of linker histones. Moreover, H1-bound states are shown to have increased stiffness within tetra-nucleosomes, but not between them. This increased stiffness leads to stronger long-range correlations within the fiber, which may result in the propagation of epigenetic signals over longer spatial ranges. These simulations highlight the effects of linker histone binding on the internal dynamics and global structure of poly-nucleosome arrays, while providing physical insight into a mechanism of chromatin compaction.

**Significance:** Linker histones dynamically bind to DNA in chromatin fibers and serve as epigentic regulators. However, the extent to which they influence the gamut of chromatin architecture is still not well understood. Using molecular dynamics simulations, we studied compact octa-nucleosome arrays with and without the H1 linker histone to better understand the mechanisms dictating the structure of the chromatin fiber. Inclusion of H1 results in stabilization of the compact chromatin structure, while its removal results in a major conformational change towards an untwisted ladder-like state. The increased rigidity and correlations within the H1-bound array suggests that H1-saturated chromatin fibers are better suited to transferring long-range epigentic information.

## Introduction

Serving as the primary storage vessel of genomic information within eukaryotic organisms, chromosomes consist predominantly of organized, long condensed fibers of DNA and structural proteins.^1^ Known as chromatin, these fibers are made of compacted repeating arrays of distinct DNA–protein complexes called nucleosomes.^1–4^ Nucleosomes consist of ~147 bp of DNA wrapped around an octamer core of organized histone proteins.^5^ Within the array, nucleosomes are inter-spaced between varying lengths of linker DNA, which is often quantified by their nucleosome-repeat-length (NRL).^6^ Computational modeling and topological studies have shown that the NRL regularity can directly affect chromatin compaction via variations in local fiber stiffness.^7,8^ Furthermore, this value can depend on interactions with a variety of cosolute compounds, nucleosome remodeling factors,^6^ or DNA sequence^9^ usually related to a level of charge neutralization and/or structural accommodation. Some examples include cosolute cations (i.e. Mg^2+^, nuclear polyamines, etc.), basic amino acids found on the terminal tail domains of core histones, proteins found outside of the nucleosome core, and H1 linker histones.^10–15^

Structural studies have shown chromatin fibers adopt multiple states, including solenoid^16,17^ or zigzag^18–23^ like-conformations, with evidence of both forms being present within the same fiber.^24^ At high ionic strength, nucleosome arrays compact to create fibers with a diameter of about 30-nm in a closed zigzag conformation,^22,25–29^ similar to what is shown in Figure 1. However, despite the fact that canonical chromatin does form chains with regular and irregular zigzag structures, there is a particular absence of 30-nm fibers from eukaryotic nuclei,^30–37^ except within terminally differentiated cells.^38–41^

**Figure 1:**
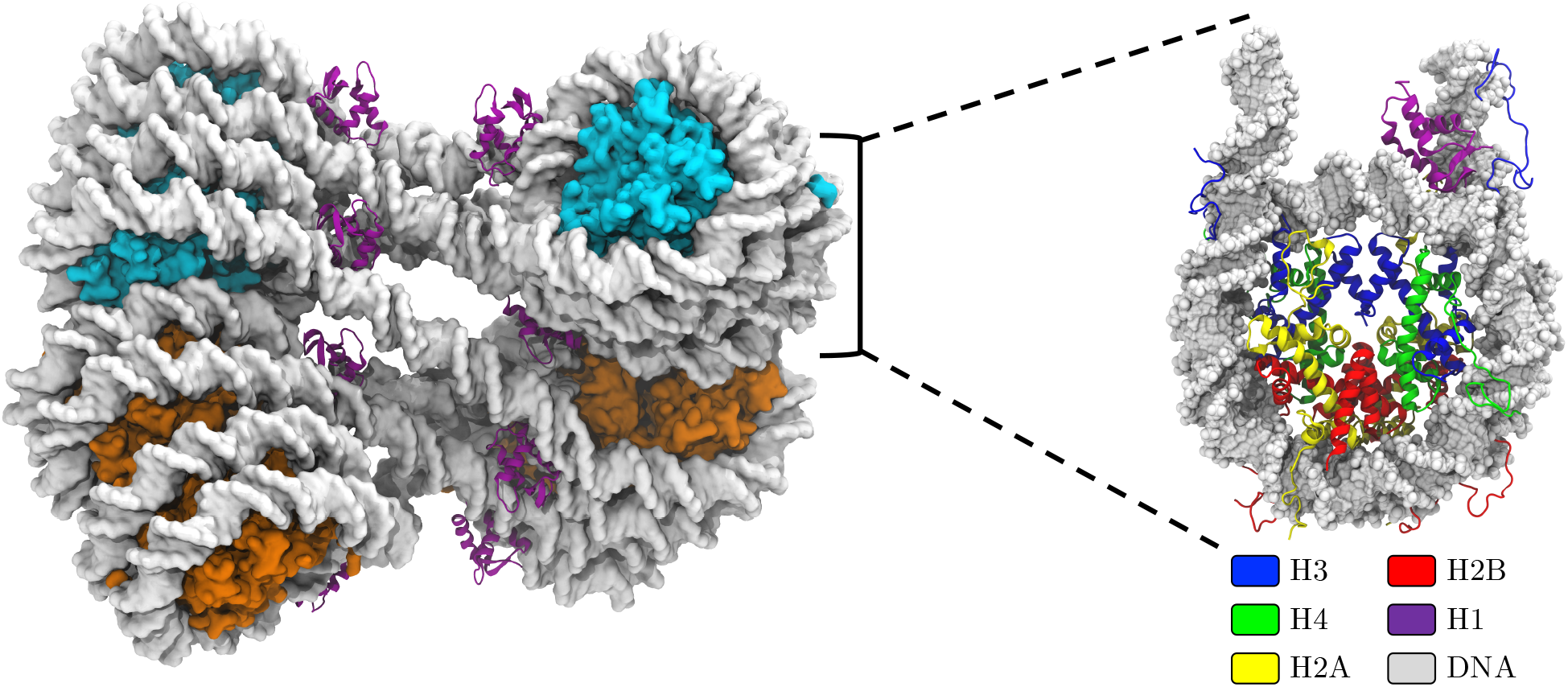
Shown on the left is the octa-nucleosome array constructed using the cryo-EM map of the 30 nm chromatin fiber.^22^ Each nucleosome is paired with a linker histone (purple) bound asymmetrically off the dyad axis. On the right is an example of a mono-nucleosome unit from the array with each individual histone shown. The core histones in the poly-nucleosome (left) are colored cyan and orange to distinguish between the upper and lower tetra-nucleosome sub-units.

To date, chromatin structural and mechanistic studies have largely focused on structural regulation at the single-nucleosomal level, ^42^ including such phenomena as nucleosome opening,^43–45^ the influence of extra-nucleosomal proteins,^46–48^ and histone variants. ^49^ More recently, studies involving poly-nucleosomal arrays and models of higher order structures have begun to show that chromatin exists in a dynamic equilibrium of states,^24,33,37,50–52^ suggesting that it exhibits large-scale, concerted dynamics orchestrated by motifs such as remodeling factors and histone variants. Moreover, contemporary coarse-grained modeling of poly-nucleosome arrays with H1 have further emphasized the diversity of chromatin dynamics highlighting structures with irregular NRLs, ^50^ varied cation concentrations,^24,51^ and higher order structures.^37,52^ However, there is a severe lack of atomistic resolution studies of poly-nucleosome arrays^53,54^ which may provide residue-specific information otherwise lost by coarse-grained models. Interacting with both core and linker DNA, ^10^ the linker histone (H1) plays a crucial role in the condensation of nucleosome chains into higher order architecture,^48,55–57^ like the zig-zag structure, ^37^ along with other cellular functions^57^ such as gene expression,^58,59^ heterochromatin genetic activity,^60^ and cell differentiation,^61,62^ among many others.^63–65^ They are found roughly every 200 ± 40 base pairs,^66^ but may be spaced more intermittently to regulate DNA accessibility for transcription factors. Additionally, linker histones predominantly interact electrostatically with the backbone phosphates of DNA using positively charged residues,^67–69^ which stabilizes nucleosome arrays hindering linker DNA accessibility and competing with core histone tails for binding space.^26,48,70–73^ However, this effect has been shown to be completely abrogated upon the addition of nucleosome-free regions within H1-saturated arrays. ^74^

In a previous study, we used all-atom molecular dynamics (MD) simulations to demonstrate that the linker histone binding mode on nucleosomes can have substantial effects on linker DNA dynamics. ^75^ Furthermore, we postulated that its presence would have cascading effects on higher order chromatin structures. Indeed, this is highlighted in many of the previously mentioned studies, but none of which provide a mechanism detailing the atomistic dynamics of the chromatin fiber in and out of the presence of H1. To extend these ideas to the chromatin fiber, we examined these dynamics via all-atom MD simulations of an octa-nucleosome array with and without the *D. melanogaster* generic globular domain of H1 bound asymmetrically off the dyad. Results suggest that linker histones provide stabilization to the fiber structure at multiple levels. Helical parameters, inspired by similar DNA base-pairing metrics,^76–78^ quantified a major conformational shift from a twisted condensed state to an untwisted ladder-like state. Stiffness parameters of these metrics show H1 binding increases torsional stress within tetra-nucleosome sub-units. Furthermore, while an angular analysis of linker DNA motions shows that linker histones limit sampling, it also highlights the stark contrast in mono-versus poly-nucleosome dynamics, especially among in-plane DNA motions. Moreover, generalized correlation analyses shows that linker histone saturation strengthens long-range correlations throughout each system, which can lead to further transfer of epigenetic information across the fiber. This linker histone saturation provides stabilization to the highly strained linker DNA resulting in a highly compact system that is unfavorable for transcription factor binding. Complete abrogation of these extra-nucleosomal proteins allows the fiber to untwist and thus alleviating the aforementioned linker DNA strain.

## Results

### Linker Histones Stabilize Tetra-Nucleosome Repeats

Models of compact octa-nucleosome structures were generated through a combination of manual placement and flexible fitting of the 1KX5 nucleosome^79^ and H1 linker histone crystal structures^75^ into the cryo-EM map by Song *et al.* (see Methods).^22^ To quantify the configurations of these complexes, we took advantage of their double-helical like structures and measured the six canonical parameters of rise, twist, roll, tilt, shift, and slide, which are typically associated with DNA basepairs (see Methods for definitions). For each system there are three sets of tetra-nucleosomal sections, which for clarity we refer to as the top, middle, and bottom segments of the array and that contain nucleosomes one through four, three through six, and five through eight respectively (Figures 2 and 3). Over our simulations, most of these metrics maintained values close to zero, with the exception of the inter-nucleosomal rise and twist which largely describe the observed large-scale conformational changes.

**Figure 2:**
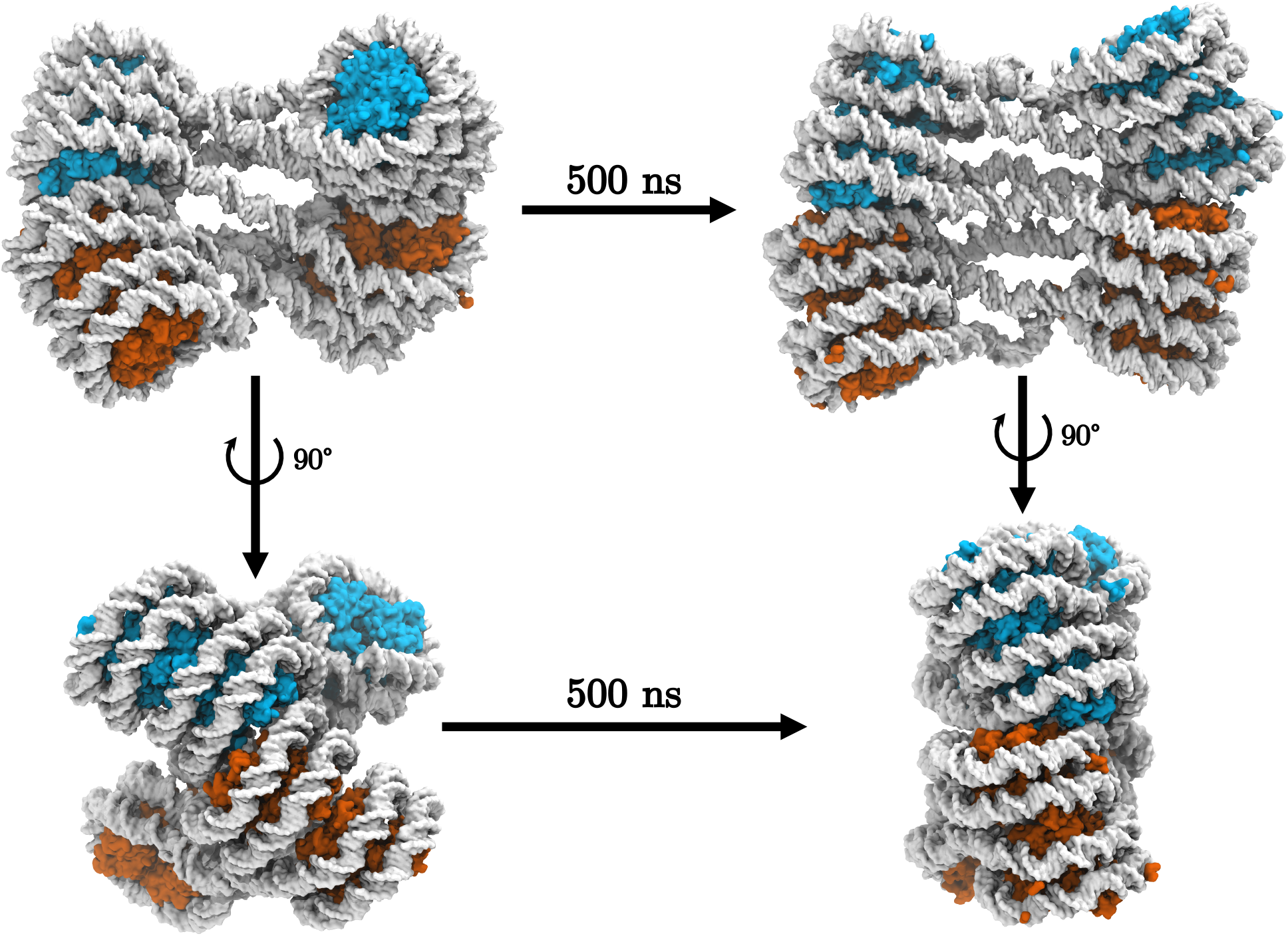
Shown are selected images from a system without the linker histone, at the beginning a simulation (left) and after 500 ns of production (right). The core histones in the poly-nucleosomes are colored to represent the different tetra-nucleosome sub-units as in Figure 1.

**Figure 3:**
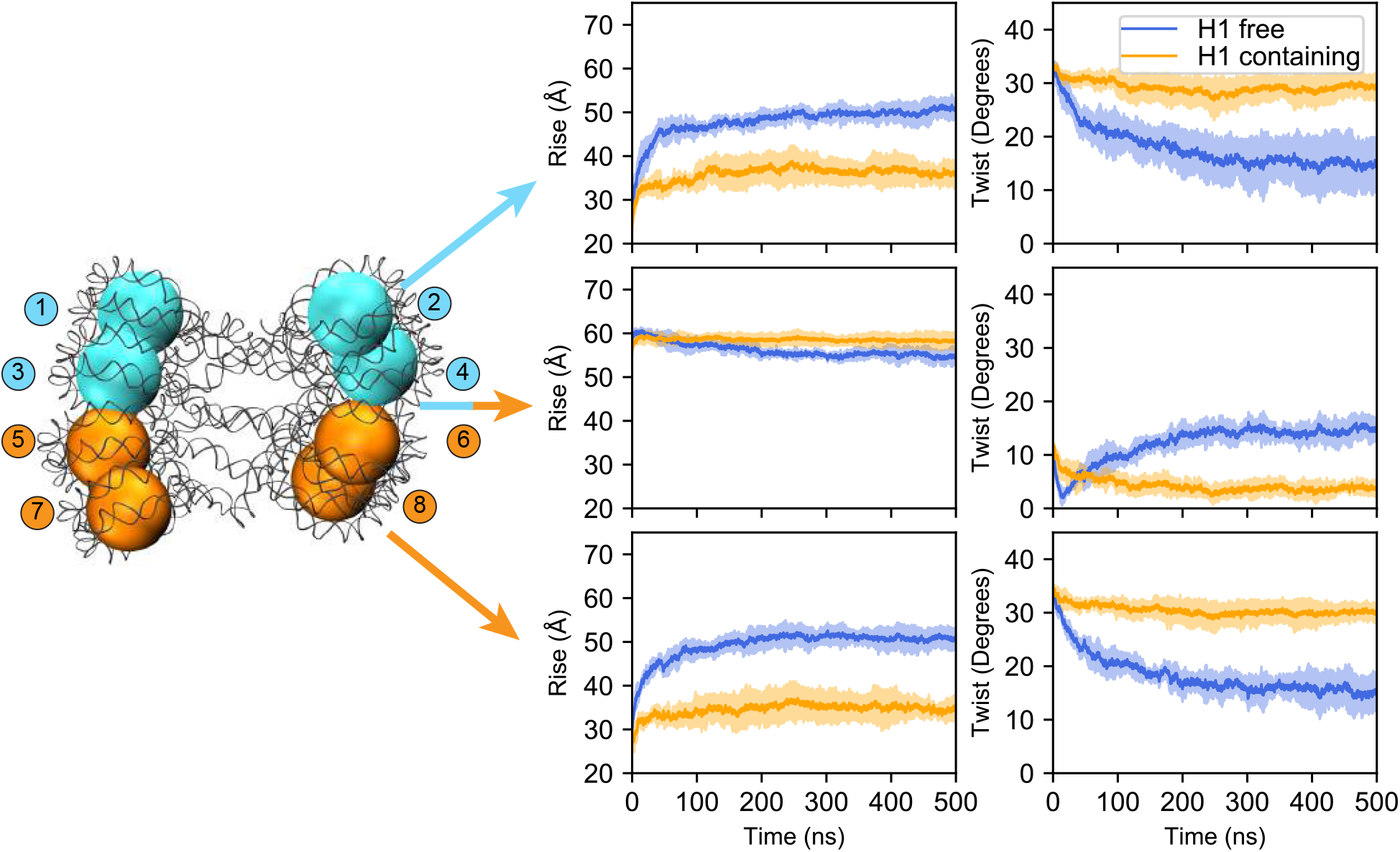
Nucleosomal rise and twist during simulations between the top, middle, and bottom four nucleosomes. Simulations with H1 (orange) maintain the initial stacked tetranucleosome structure, whereas simulations without H1 (blue) adopt a looser stacked conformation. Shown are the average and standard deviations (shaded regions) between the three simulations for each system. Colors of the octa-nucleosome array are to distinguish between tetra-nucleosome sub-units such as in Figures 1 and 2. Each nucleosome is designated with a number from 1 to 8 which is referenced throughout this manuscript.

In each of the three 500 ns simulations we performed with H1, the rise and twist parameters remained similar to their initial values (middle and right of Figures 2, 3, and S1-S2). In particular, for both the bottom and top tetra-nucleosome segments (tetraNuc), the initial rise and twist of ~27 Å and 33° were largely maintained, equilibrating at ~5 Å and 30°. In contrast, for the middle segment the initial and final rises were higher, with an average value of 58 Å, with a reduced twist that equilibrated from 14° to 4°. This difference in values for the top and bottom array segments relative to the middle highlights the difference in the intra- and inter-tetraNuc structures: in a tetraNuc unit there is relatively little rise between nucleosomes (Nuc), as Nuc_*i*_ forms a tight packing interface with Nuc_*i*+2_ that creates a twist around the central fiber axis. Meanwhile, between tetraNuc structures the inter-nucleosomal packing is reduced and there is looser interface that has a higher rise and less twist around the helical axis. In addition, the minimal changes in these parameters during each simulation, and their reproducibility between each independent simulation (Figures S1-S2) demonstrates the stability on this stacked tetraNuc structure on the hundreds of nanoseconds timescales.

In contrast, in each of the three 500 ns simulations without H1, there were dramatic and irreversible changes in all measured rise and twist values which resulted in an elongated and less twisted array (Figures 2 and S6). Despite starting with values identical to H1 arrays, the stacked tetra-nucleosome structure was lost in the first 150 ns, as is evidenced by the increase in rise of the top and bottom sections to 51 Å and the decrease in twist to 15°. These values approach those of the middle array segment, which equilibrate to 53 Å with an identical twist of 15°. This close agreement between the rise and twist for the middle with the top and bottom array segments demonstrates that the stacked tetraNuc structure is lost, as there is little physical difference between the structures of nucleosomes 1-4 and 5-8 with 3-6. This large conformational change is also demonstrated by the elevated root mean square deviation (RMSD) values for the C_α_ and phosphate atoms, which ranged between 39 and 47 Å for H1 lacking arrays, relative to a range of 15-18 Å for H1 containing arrays (Figure S3).

### Greater Poly-Nucleosome Architecture Dictates Linker DNA Sampling

Linker histones have a direct effect on the motions of linker DNA and nucleosomal DNA through favorable energetic interactions driven by electrostatic and Van der Waals forces. In a previous study, we emphasized the significance of these interactions by demonstrating how the linker histone binding pose, along with how their mere presence, can affect experimental results. ^75^ Here, we translated these ideas to the context of poly-nucleosomal arrays by plotting the in- and out-of-nucleosomal-plane motions of both linker DNA arms in Figure 4 and Supplementary Figure S7, respectively. In general, removing linker histones from the array not only results in overall increased DNA sampling, but the development of novel linker DNA states. This is especially evident in the terminal nucleosomes, labelled Nuc 1, Nuc 2, Nuc 7, and Nuc 8 in Figures 4 and S7. Nucleosomes 2 and 7 presented the most structural distortion, which can be attributed to the drastic global change in conformation which occurred in the all simulations lacking linker histones.

**Figure 4:**
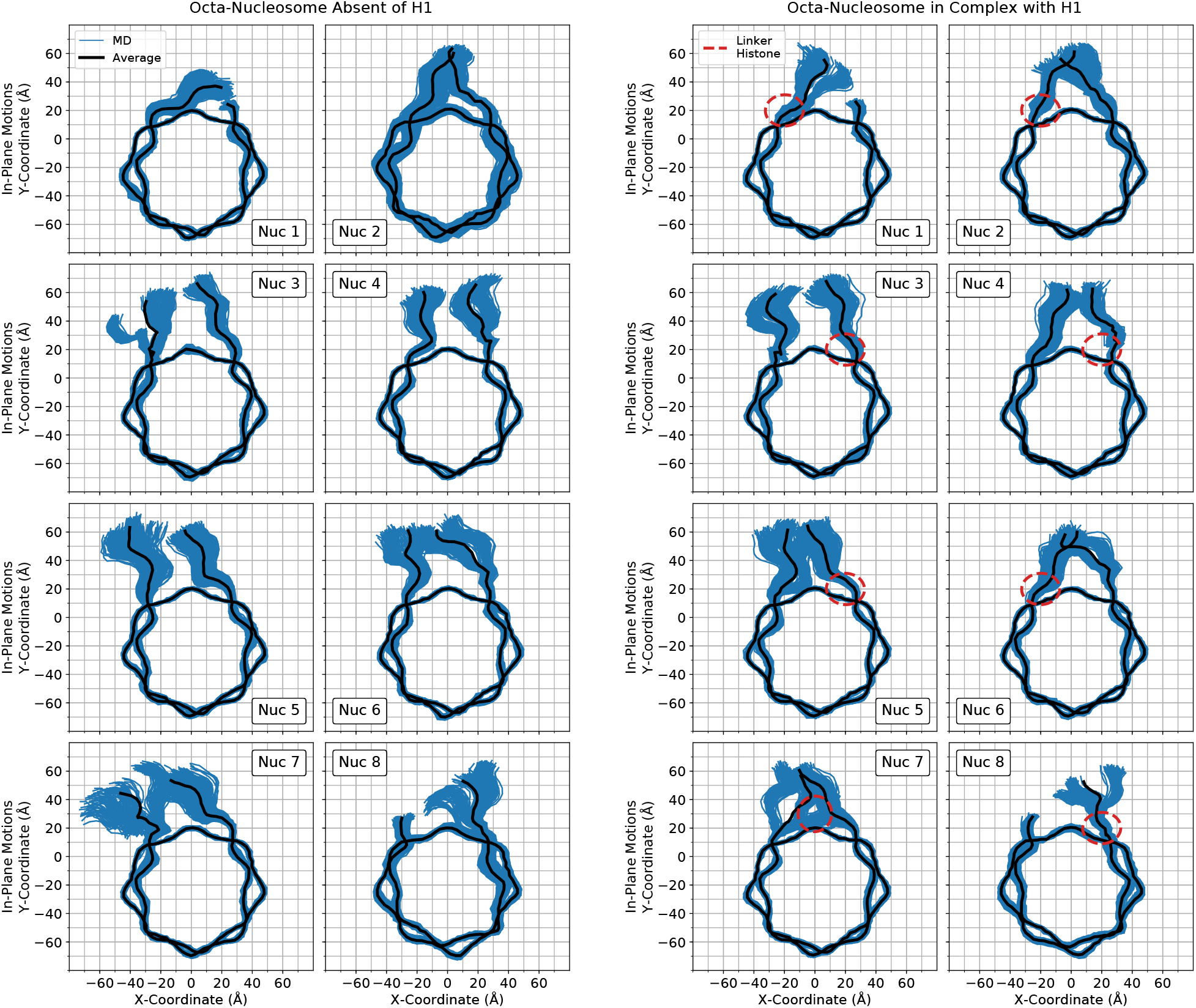
In-plane (top) DNA motions sampled by the octa-nucleosome arrays absent (left) and in complex (right) with the linker histone H1. Shown in blue are configurations sampled throughout the MD simulation (263 representative frames - every 4 ns of simulation time) while the average configuration is shown in black. For reference, the relative position of each nucleosome in its array is labeled in the corner of each graph. This label is consistent with the numbering in Figure 3. Additionally, the approximate position of the linker histone is shown as a dashed-line red ellipse. Figures inspired by work from Shaytan *et al*^80^ and single comparative nucleosome results were published previously. ^75^

To quantify these dynamics, the in- and out-of plane linker DNA motions were calculated and denoted as the α- and β-angles, respectively (as inspired by Bednar *et al.,* see Methods and Supporting Information for detailed definitions, Figures 5 and S4). The α-angles relate predominately to fluctuations in DNA breathing and ranged from −112.2° to 96.9° with an average of 35.4°. Out-of-plane motions, or β-angles, ranged from −56.4° to 62.9° and averaged 1.1°. An additional observation was that the entry and exit DNA arms of H1-absent nucleosomes had somewhat different probability distributions, which can be attributed to the asymmetric initial conformation within the poly-nucleosome array and the asymmetric nucleic acid sequence of Widom 601. While we previously illustrated that linker histone binding alters linker DNA fluctuations, Figure 5 shows that those effects are more pronounced in poly-nucleosome systems. For example, the α-Entry angle sampling range was reduced by 56.7° when the octa-nucleosome is saturated with H1. However, this large reduction in sampling space is not present in the α-Exit angles. This occurs because the majority of the structural strain within the compact array is distributed onto the Entry linker DNA. When the linker histone is no loner present, the distorted Entry DNA must endure the bulk of the conformational alleviation. In contrast to the mono-nucleosomes, the octa-nucleosome arrays sample a wider breadth of angles, particularly the α-angle dimension. However, the individual nucleosomes that constitute the array do not readily transition throughout this entire phase space. More often, it is the case that each nucleosome will exist as a single stable, but independent, state unable to sample much outside of its respective potential well. Despite the apparent increased sampling of angles within the array, there is an inherent entropic cost for each individual nucleosome as oppose to existing solitary in solution.

**Figure 5:**
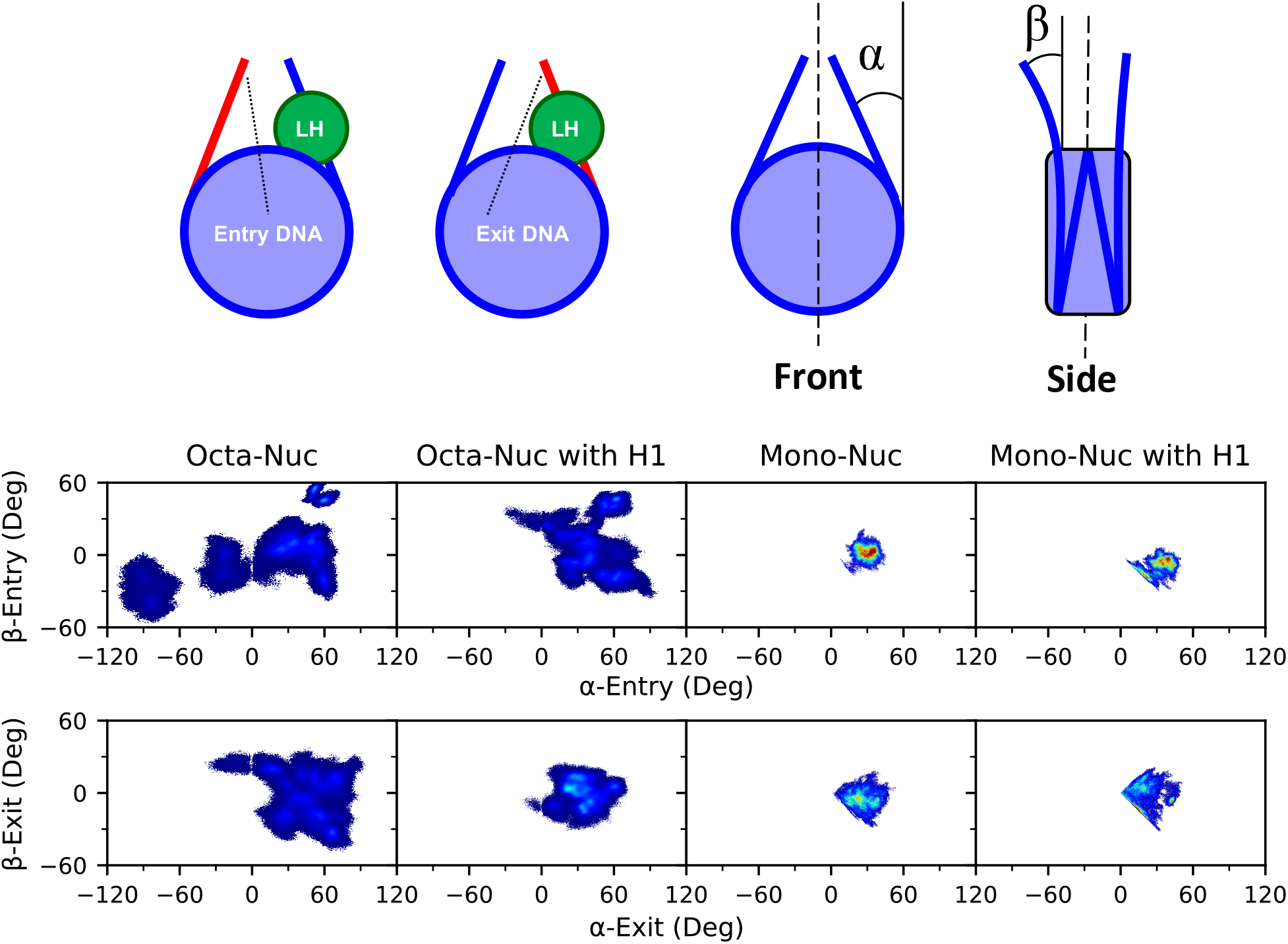
Comparison of DNA sampling for the entry (top plots) and exit (bottom plots) DNA segments for systems with and without H1. The α-angles described in-plane motions, whereas β-angles describe out- of-plane, as depicted in the diagrams on the to of the figure. For clarity, the entry- and exit-DNA segments are depicted in the top left diagram with the linker histone (LH), if present, in green. Density is represented as a gradient from blue (low density) to red (high density). Mono-nucleosome results are from previously published results. ^75^

Although mono-nucleosomes had more available conformational freedom, linker DNA in poly-nucleosomes sample a broader spectrum of states throughout the simulations, which we attribute to the strained nature of nucleosome arrays biasing linker DNA into ordinarily unattainable states, as observed in Figure 4. To further quantify these differences, we calculated Jensen-Shannon distances (JS_*dist*_) (Table 1), based on the Jensen-Shannon divergence (JSD), between the one-dimensional probability distributions displayed in Figure S4. Values closer to zero correspond to a greater similarity in probability distributions, whereas values approaching 1.0 correspond to a greater dissimilarity. The JS_*dist*_ values generally exhibit a stark contrast between mono-nucleosome and octa-nucleosome systems with values often above 0.50, although with some exceptions. In particular, α/β-angles of the Exit DNA (in contact with H1) sample a much more similar phase space than the Entry DNA angles. Additionally, highlighted by a JS_*dist*_ of 0.72, binding of the H1 severely alters sampling of β-Entry angles in mono-nucleosomes and thus presenting an extreme case for which to compare poly-nucleosome systems.

**Table 1:** Jensen-Shannon distances (equation (4)) for one dimensional probability distributions (Figure S5) of DNA motions between systems. For clarity, comparisons with low differences (JS_*dist*_<0.20) are in blue, increased differences (0.20<JS_*dist*_<0.40) are in green, high differences (0.40 <JS_*dist*_<0.60) are in orange, and very high differences (0.60<JS_*dist*_<1.00) in red. The lower numerical values correspond to a greater similarity in probability distributions, whereas higher numerical values correspond to a greater dissimilarity. Two identical distributions will produce a Jensen-Shannon distance of 0.00, whereas distributions that do no share any phase space commonality will result in 1.00.

| | α-Entry | | | α-Exit | | |
| --- | --- | --- | --- | --- | --- | --- |
|  | MonoNuc<br>with H1 | OctaNuc | OctaNuc<br>with H1 | MonoNuc<br>with H1 | OctaNuc | OctaNuc<br>with H1 |
| MonoNuc | 0.35 | 0.69 | 0.64 | 0.35 | 0.62 | 0.36 |
| MonoNuc with H1 |  | 0.63 | 0.54 |  | 0.67 | 0.54 |
| OctaNuc |  |  | 0.42 |  |  | 0.41 |

| | β-Entry | | | β-Exit | | |
| --- | --- | --- | --- | --- | --- | --- |
|  | MonoNuc<br>with H1 | OctaNuc | OctaNuc<br>with H1 | MonoNuc<br>with H1 | OctaNuc | OctaNuc<br>with H1 |
| MonoNuc | 0.72 | 0.53 | 0.64 | 0.30 | 0.59 | 0.36 |
| MonoNuc with H1 |  | 0.69 | 0.58 |  | 0.44 | 0.15 |
| OctaNuc |  |  | 0.39 |  |  | 0.46 |

### Linker Histones Lead to Stiffer Nucleosome Arrays

To characterize the effects of linker histones on the flexibility of compact nucleosomal arrays, the local elastic properties of these systems were computed based on the helical parameter covariance matrices (see Methods for details). For each of the diagonal elements in these matrices, the associated force constants were equal or higher for H1 containing arrays as compared to H1 free arrays (Figure 6). However, in many cases the stiffness, and the difference between the H1 free and containing systems, was dependent on whether inter- or intra-tetra-nucleosome units were considered. For example, in consideration of the rise parameter, in H1-free arrays all of these elastic constants had values that were approximately equal to one another (within the standard error). In H1-containing arrays the stiffness parameters were similar, however, given the smaller standard errors we are able to conclude that the inter-nucleosome rise stiffness is sightly higher than the intra-nucleosome stiffness. Other parameters, such as slide shift, tilt, and roll, displayed a similar trend that any difference between the stiffness parameters were small, and close to the standard errors. In contrast, linker histones created a significant increase in the stiffness of the twist within tetra-nucleosome units, but did not influence the force constants between them, suggesting that stacked tetra-nucleosome units impart resistance in poly-nucleosomal arrays via torsional stress.

**Figure 6:**
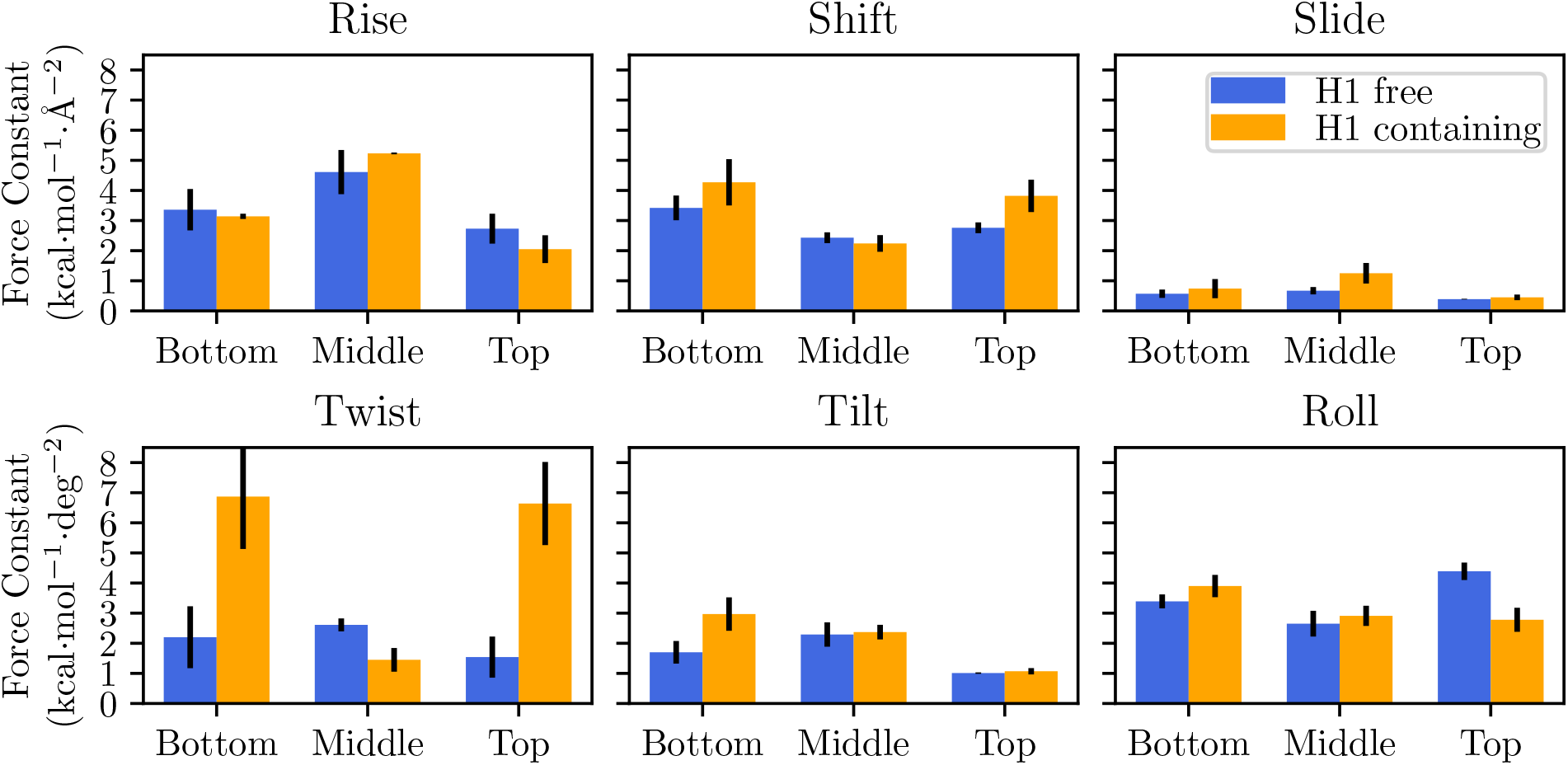
Force constants of helical parameters for the Bottom, Middle, and Top tetra-nucleosomal structures (as defined in Figure S1) and reported in Tables S1 and S2. Error bars represent the standard error of the mean computed from all three simulations.

While the on-diagonal elements of the stiffness matrices are the force constants for the canonical helical parameter, the off-diagonal elements correspond to the coupling between these parameters (Tables S1-S2). The majority of these elements are small and within the standard error of zero, indicating that these degrees of freedom are largely uncorrelated from one another. In contrast, the twist-rise coupling is significant and shows a pattern similar to the twist force constants: without H1 the coupling constants range from 1.60 ± 0.51 to 2.95 ± 0.24 kcal·mol^-1^ ·Å^-1^ ·deg^-1^, with the highest being for the middle segment of the octa-nucleosome array. In contrast, there is a more significant difference with H1, where the bottom and top tetra-nucleosome segments contain coupling constants of 3.92 ± 0.45 and 3.08 ± 0.69 kcal·mol^-1^ ·Å^-1^ ·deg^-1^, and the middle has a significantly reduced value of 1.21 ± 0.24 kcal·mol^-1^ ·Å^-1^ ·deg^-1^. This further points to the increased rigidity within linker histone stabilized tetra-nucleosome units, and the relative looseness in these arrays between them. In addition, the positive values observed for all twist-rise coupling constants show that these arrays contract upon overtwisting, which is in line with what one would expect from models of simple helical elastic polymers but is contrary to DNA which elongates when overtwisted.^81,82^

### Linker Histones Create Long Range Correlations

Having established that linker histones create stiffer nucleosomal arrays, we sought to understand the implications for larger scale dynamical properties. We therefore performed a generalized correlation analysis on each system to determine the pairwise correlations between each DNA base and protein residue in the system.^83^ The average H1 free correlation matrix shows the expected behavior that within each nucleosome there is a high correlation, as individual nucleosomes are highly rigid on the nanosecond timescale (see red square in Figure 7a). In addition, core histones are highly correlated with the DNA which wraps around it, as evidenced by the red patterns at the top and right side of these figures, and DNA bases are highly correlated with the base they are paired with, as shown by the “X” mark in the upper right hand corner. More interestingly, individual nucleosomes are highly correlated with nucleosomes that stack directly above or below them. That is, Nuc_*i*_ is highly correlated with Nuc_*i*+2_ and Nuc_*i*−2_. In contrast, nucleosomes did not have a large correlation with their adjacent nucleosome, as the Nuc_*i*_ and Nuc_*i*+1_ correlations were relatively low.

**Figure 7:**
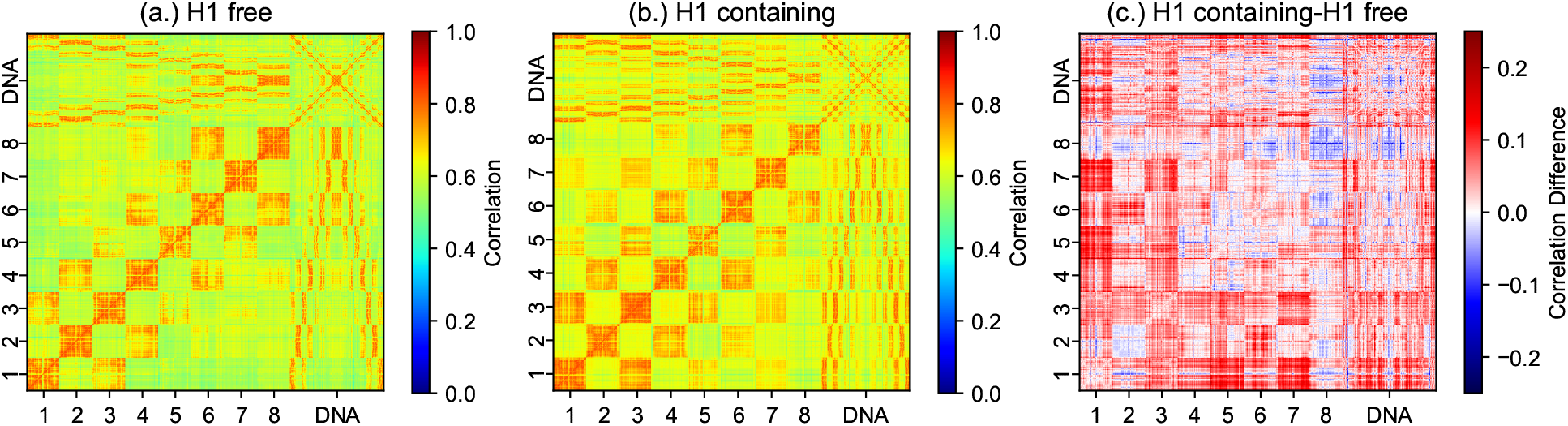
Inter-residue correlations for systems lacking (a.) and containing (b.) H1. H1 increases system correlations, notably through increased correlations in stacked nucleosomes, as shown in the difference between system with and without H1 (c.).

Upon the addition of linker histones, the overall pattern of strong intra-nucleosomal and local nucleosome/DNA correlations remained (Figure 7b). In addition, the majority of correlations were enhanced, as evidenced in the difference map between the correlations in the H1 containing and free systems (Figure 7c). Of particular interest are the stronger correlations between all of the odd numbered and even numbered nucleosomes, that is between nucleosomes 1,3,5, and 7 and 2,4,6, and 8. This suggests that in the more compact and stiffer arrays induced by linker histones, correlations are able to propagate throughout each side of these two-start zig-zag arrays much further then they can without linker histones.

## Discussion

Here, we have used conventional MD simulations to probe the effects of linker histone binding on an octa-nucleosome model of the chromatin fiber. We calculated nucleosomal helical parameters quantifying a global conformational shift that demonstrate the importance of H1 in the structural stability of chromatin fibers. Moreover, we have captured a physical phenomenon that has been rarely observed experimentally^51^ - the helical untwisting of the poly-nucloeosme array. The most probable explanation for this occurrence is the lack of strong protein-DNA interactions provided by the linker-histone/DNA motif. The chromatin fiber is highly compact and very strained by our observations. The saturated binding of linker histones to the linker DNA stabilizes the system and prevents it from untwisting. Presumably, this untwisting effect could be mimicked by a decrease in salt concentration. In fact, Garcia-Saez *et al.* presented an untwisted nucleosome array model based on cryo-EM data of arrays under low-salt conditions. Interestingly, this phenomenon was not hindered by the presence of the linker histone, but occurs readily in saturated arrays. One potential explanation for this discrepancy is that the Garcia-Saez *et al.* poly-nucleosome array exhibited a low packing density resulting in inherently less inter-nucleosome protein-DNA interactions. This can be attributed to longer linker DNA length between nucleosomes (50 bp versus 30 bp) and the more rigid on-dyad binding mode. As shown in Figures 4 and 5, our more compact model requires more diverse sampling of the linker DNA, which was previously shown to be hindered upon on-dyad binding. ^75^ A compact chromatin fiber with H1 bound on-dyad would be more strained than its off-dyad bound counterpart and easily perturbed upon a reduction in ionic strength, as observed by Garcia-Saez *et al*.

By altering linker DNA dynamics, linker histones inherently inhibit transcription and promote the compaction of chromatin fibers. Our simulations have shown how substantial an impact the linker histones have on the global chromatin compaction. This effect is highlighted by the dramatic reduction in sampling space upon H1 binding. Once bound, the linker histone increases the rigidity of the entire system, as was further emphasized by the increase in correlation throughout the fiber (Figure 7). Furthermore, the disparity in sampling between mono- and poly-nucleosomes is extensive. We accredit this observation to the unique structure of the chromatin fiber. In mono-nucleosomes, linker DNA is generally free to move about in solution, unless bound to a linker histone. However, in poly-nucleosome arrays each nucleosome must adopt a specific conformation to alleviate strain on the entire system. Here, we stress caution when studying mono-nucleosomes and deducing conclusions about their dynamics. The Jensen-Shannon distances in Table 1 highlights the vast dissimilarities between the two systems, specifically with in-plane linker DNA motions, and why results from mono-nucleosome studies, especially related to DNA, may not be transferable when describing the greater chromatin architecture.

The octa-nucleosome system studied here is composed of two distinct, although attached, tetra-nucleosome sub-units, as illustrated throughout this manuscript by light-blue and orange colored core histones (Figures 1, 3, 2, and S6). Using force constants derived from the helical parameters, we found that linker histones impart increased torsional stress within these tetra-nucleosome units while slightly decreasing it between units. Interestingly, linker histones between tetra-nucleosome sub-units are quite close proximity to one another and have been speculated to interact,^22,51^ giving rise to potential sites for post-translational modifications.^84–87^ Our calculations show only a few inter-linker histone contacts, specifically between linker histones associated with Nuc_4_ and Nuc_6_ (Figures S8 and S9). Unfortunately, the presence of these interactions did not contribute to the overall stiffness of our model. This is evident by the aforementioned force constants which show a decrease in torsional stress between tetra-nucleosome sub-units upon H1 binding. Therefore, by our observations, inter-linker histone interactions do not significantly contribute to chromatin compaction in the octa-nucleosome model studied here. However, it should be stated that this model included only the globular domain of each linker histone and not the N- and C-terminal tails, known to interact with H3 tails to facilitate binding, ^88^ translating to an examination of localized interactions. The inclusion of these motifs may lead to more interactions with other linker histones, linker DNA, and/or nucleosomes, resulting in increased chromatin fiber compaction and rigidity.

In a recent comprehensive study, Perišić *et al.* used meso-scale modeling to demonstrate the extent to which linker histone binding modes and variants affect chromatin compaction. ^89^ They were able to connect existing ideas suggesting that combinations of on- and off-dyad binding result in varying levels of compaction on a spectrum between condensed^22^ and uncondensed^90^ arrays, respectively. Their work provided strong reinforcement that shifts between these two states are directly associated with a shift in linker histone binding mode, ^51^ a sentiment which we share. ^75^ Here, we quantified the effects of linker histones on condensation using various metrics from an atomistic perspective. Furthermore, this work demonstrates that chromatin can experience large conformational transitions in timescales of under a microsecond, which is well under the time it may take to expose nucleosomes for transcription and DNA repair.^91–93^

## Materials and Methods

### System Construction

Core histones and the asymmetric Widom 601 DNA were modelled based on the 1KX5 crystal structure (resolution 1.94 Å ^79^). Missing residues and nucleotides were added using Modeller via the Chimera graphical user interface.^94,95^ Nucleosomes were manually placed to fit within the 12 Å cryo-EM map,^22^ followed by 30 bp of linker DNA between them, which was built using the Nucleic Acids Builder (NAB) module within AmberTools18 software package.^96^ From here, rigid docking was performed using the Colores module of Situs,^97,98^ which was followed by flexible docking using internal coordinates normal mode analysis (iMOD).^99^ Finally, we built, placed, and validated the linker histone binding mode within the octa-nuclesome array using methods detailed in our previous work. ^75^ When solvated, these systems were approximately 2,757,000 atoms.

### Molecular Dynamics simulations

All systems were prepared and simulated using the GROMACS 2016.4 software package.^100^ Each system was solvated in a TIP3P water box extending at least 10 Å from the solute.^101,102^ Using Joung-Cheatham ions,^103,104^ the solvent contained 150 mM NaCl, sodium cations to neutralize negative charges, and magnesium ions that replaced the manganese ions in the 1KX5 crystal structure. Only magnesium ions in the DNA grooves were included, whereas those located close to the the linker histones binding locations were excluded so as to not interfere with LH-DNA interactions. The AMBER14SB and BSC1 force fields were used for protein and DNA interactions, respectively.^105,106^ A cutoff distance of 10.0 Å with a switching function beginning at 8.0 Å was used for nonbonded interactions, and long range electrostatics were treated with particle mesh Ewald calculations.^107^ Systems were minimized for 10,000 steps, and then equilibrated for 100 ps at constant volume and temperature and for 1 ns at constant pressure and temperature. Production simulations were carried out for 500 ns in the NPT ensemble, using a Parrinello-Rahman barostat^108^ with a time constant of 1.0 ps to control the pressure and a Nosé-Hoover thermostat at 300K with a time constant 0.5 ps. Electrostatic interactions were treated with the Particle-Mesh Ewald (PME) method^107^ and 10 Å cut-off. Simulations were conducted on two systems: one with the linker histone bound to the DNA of each nucleosome and one without the presence of H1. Each simulation was run in triplicate for 500 ns with a 2 fs timestep using resources provided by the Extreme Science and Engineering Discovery Environment (XSEDE).^109^

### Analysis

#### Helical Parameters

The local double helical structure of the chromatin fiber was quantified based upon three translational (rise, shift, and slide) and three rotational (twist, tilt, and roll) parameters.^77^ In analogy with DNA structure, a “base pair” was defined as two nucleosomes that were in the same z-plane with one another, where the z-axis is defined as the principal fiber axis. For each nucleosome pair, the local x-axis was defined as a vector from the center of mass of the DNA phosphate atoms of NCP *i* to the center of mass of the DNA phosphate atoms of NCP *i* + 1. The z-axis was defined as the average of the third principal axis of inertia of NCPs *i* and *i* + 1, as computed with gromacs, and the y-axis was computed as the vector perpendicular to these two vectors. Following the definition of these “base-pair” axis, the algorithm outlined by Lu *et al.* was used to compute the rise, shift, slide, twist, tilt, and roll. ^76^

To compute the elastic force constants, a harmonic approximation was made for each of the basepair parameters, such that the internal energy of the fiber is estimated by:

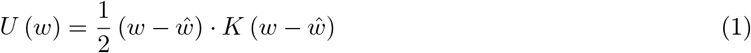

where *w* is the vector of inter-base pair parameters, 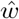 is their mean, and *K* a stiffness matrix.^110^ Given small fluctuations, the equipartion theorem can be used to construct K from the inverse of the covariance matrix, C:

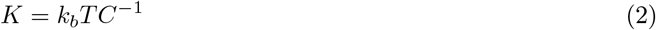

where *k_b_* is Boltzmann’s constant, and T the system temperature. Terms along the diagonal of K are the individual parameter force constants, and off-digonal terms represent the parameter coupling constants. Error bars were computed by calculating the stiffness matrices for each of the three simulations and taking the standard error of the mean.

#### Generalized Correlation

Mutual-information based generalized correlation methods were employed to capture non-collinear correlations between residue pairs to describe both linear and non-linear coupled motions. Results were computed using the g_correlation plugin for GROMACS/3.3.^83,100,111^ The first 150 ns of simulation time was allotted for equilibration while trajectories were analyzed every 50 ps. Analyses were performed only on the protein *C_α_* and the nucleotide C1’ heavy atoms.

#### Linker DNA Dynamics

The linker DNA in- and out-of nucleosomal plane motions were quantified to describe the linker DNA motions. To define the plane, the nucleosomal DNA was divided into four quadrants and the center of mass of the C1’ atoms within the two quadrants located distal from the linker DNA were used for two points, while the third point was defined as the C1’ center of mass of bases 83 and 250 which are located approximately on the dyad axis (see previous work^75^ for details). The linker DNA vectors were defined as the C1’ center of mass of the base pairs at the origin of the linker DNA (bases 20-315 and 148-187) and terminal base pairs (bases 1-354 and 177-178), respectively. The α-angles were defined as in-plane and the β-angles were defined as out-of-plane motions of this vector. Positive α-angles were defined as inward motions towards the dyad axis while positive β-angles were defined as outward motions away from the nucleosomal-plane. For reference, the angles shown in Figure 5 are positive.

The change in linker DNA sampling between H1-bound and -free systems were computed using two metrics, the Kullback-Leibler^112^ (KLD) and Jensen-Shannon divergences (JSD),^113,114^ respectively:

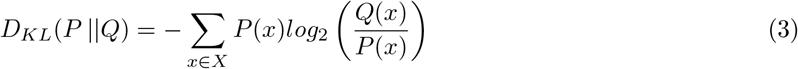

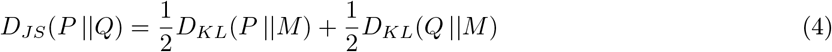

where,

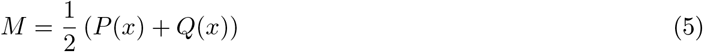

where, in equation (4), *Q*(*x*) is the normalized reference distribution and *P*(*x*) is the normalized data set. In equation (4), the JSD gives equal weight to *Q*(*x*) and *P*(*x*) by calculating their KLD with respect to an average distribution, *M* in equation (5). With these measures, we are comparing two probability distributions and thus employ a base 2 logarithm as shown in equation (3). Due to its symmetric nature, the square root of the JSD can be used as a true mathematical metric known as the Jensen-Shannon distance^115–117^ which is how we have reported it in this study.

## Author Contributions

F.R.R. ran the simulations of the poly-nucleosome array. J.W. and D.C.W. ran the analyses and prepared the manuscript. All authors contributed to the design of this project.

## Acknowledgments

Work in the Wereszczynski group is funded by the National Science Foundation [CAREER-1552743] and the National Institutes of Health [1R35GM119647]. This work used the Extreme Science and Engineering Discovery Environment, which is supported by the National Science Foundation [ACI-1053575].

