## Supporting Information for "The Dynamic Influence of Linker Histone Saturation within the Poly-Nucleosome Array"

Dustin C. Woods[1] and Francisco Rodríguez-Ropero[2] and Jeff Wereszczynski[2]

[1]Department of Chemistry and the Center for the Molecular Study of Condensed Soft Matter, Illinois Institute of Technology, Chicago, IL 60616
[2]Department of Physics and the Center for the Molecular Study of Condensed Soft Matter, Illinois Institute of Technology, Chicago, IL 60616

Table S1: Stiffness matrices for octa-nucleosome structures lacking H1. Error bars are the standard error of the mean for each value as computed over three simulations. Bolded values have a standard error that is below 25% of the absolute value of the computed stiffness value.

| | | | | | | |
|---|---|---|---|---|---|---|
| H1 Free: Bottom | | | | | | |
| | Rise (Å) | Shift (Å) | Slide (Å) | Twist (Deg) | Tilt (Deg) | Roll (Deg) |
| Rise (Å) | **3.36 ± 0.69** | -0.01 ± 0.06 | -0.23 ± 0.18 | 2.38 ± 0.85 | -0.25 ± 0.18 | -0.59 ± 0.21 |
| Shift (Å) | -0.01 ± 0.06 | **3.42 ± 0.41** | -0.03 ± 0.12 | 0.15 ± 0.06 | -0.37 ± 0.26 | **-2.72 ± 0.27** |
| Slide (Å) | -0.23 ± 0.18 | -0.03 ± 0.12 | **0.57 ± 0.14** | -0.14 ± 0.06 | -0.24 ± 0.11 | 0.12 ± 0.19 |
| Twist (Deg) | 2.38 ± 0.85 | 0.15 ± 0.06 | -0.14 ± 0.06 | 2.20 ± 1.03 | -0.32 ± 0.15 | **-0.51 ± 0.11** |
| Tilt (Deg) | -0.25 ± 0.18 | -0.37 ± 0.26 | -0.24 ± 0.11 | -0.32 ± 0.15 | **1.70 ± 0.37** | 0.46 ± 0.22 |
| Roll (Deg) | -0.59 ± 0.21 | **-2.72 ± 0.27** | 0.12 ± 0.19 | **-0.51 ± 0.11** | 0.46 ± 0.22 | **3.39 ± 0.23** |
| H1 Free: Middle | | | | | | |
| | Rise (Å) | Shift (Å) | Slide (Å) | Twist (Deg) | Tilt (Deg) | Roll (Deg) |
| Rise (Å) | **4.61 ± 0.73** | -0.14 ± 0.42 | -0.22 ± 0.07 | **2.95 ± 0.24** | -0.53 ± 0.56 | 0.10 ± 0.40 |
| Shift (Å) | -0.14 ± 0.42 | **2.43 ± 0.18** | 0.21 ± 0.16 | -0.22 ± 0.15 | 0.36 ± 0.31 | **-2.25 ± 0.33** |
| Slide (Å) | -0.22 ± 0.07 | 0.21 ± 0.16 | **0.67 ± 0.12** | **-0.19 ± 0.03** | -0.25 ± 0.28 | -0.31 ± 0.09 |
| Twist (Deg) | **2.95 ± 0.24** | -0.22 ± 0.15 | **-0.19 ± 0.03** | **2.61 ± 0.21** | -0.74 ± 0.29 | 0.28 ± 0.30 |
| Tilt (Deg) | -0.53 ± 0.56 | 0.36 ± 0.31 | -0.25 ± 0.28 | -0.74 ± 0.29 | **2.29 ± 0.40** | -0.27 ± 0.34 |
| Roll (Deg) | 0.10 ± 0.40 | **-2.25 ± 0.33** | -0.31 ± 0.09 | 0.28 ± 0.30 | -0.27 ± 0.34 | **2.65 ± 0.43** |
| H1 Free: Top | | | | | | |
| | Rise (Å) | Shift (Å) | Slide (Å) | Twist (Deg) | Tilt (Deg) | Roll (Deg) |
| Rise (Å) | **2.73 ± 0.50** | 0.14 ± 0.44 | 0.07 ± 0.05 | 1.60 ± 0.51 | -0.42 ± 0.16 | -0.50 ± 0.45 |
| Shift (Å) | 0.14 ± 0.44 | **2.76 ± 0.18** | -0.05 ± 0.11 | 0.18 ± 0.33 | 0.04 ± 0.16 | **-2.98 ± 0.22** |
| Slide (Å) | 0.07 ± 0.05 | -0.05 ± 0.11 | **0.39 ± 0.01** | 0.12 ± 0.05 | 0.00 ± 0.09 | 0.08 ± 0.10 |
| Twist (Deg) | 1.60 ± 0.51 | 0.18 ± 0.33 | 0.12 ± 0.05 | 1.54 ± 0.68 | -0.39 ± 0.13 | -0.54 ± 0.40 |
| Tilt (Deg) | -0.42 ± 0.16 | 0.04 ± 0.16 | 0.00 ± 0.09 | -0.39 ± 0.13 | **1.01 ± 0.02** | 0.31 ± 0.20 |
| Roll (Deg) | -0.50 ± 0.45 | **-2.98 ± 0.22** | 0.08 ± 0.10 | -0.54 ± 0.40 | 0.31 ± 0.20 | **4.39 ± 0.29** |

Table S2: Stiffness matrices for octa-nucleosome structures containing H1. Error bars are the standard error of the mean for each value as computed over three simulations. Bolded values have a standard error that is below 25% of the absolute value of the computed stiffness value.

| | Rise (Å) | Shift (Å) | Slide (Å) | Twist (Deg) | Tilt (Deg) | Roll (Deg) |
|---|---|---|---|---|---|---|
| **H1 Containing: Bottom** | | | | | | |
| Rise (Å) | **3.14 ± 0.09** | -0.23 ± 0.20 | 0.18 ± 0.07 | **3.92 ± 0.45** | 0.09 ± 0.21 | -0.76 ± 0.23 |
| Shift (Å) | -0.23 ± 0.20 | **4.27 ± 0.77** | -0.06 ± 0.13 | 0.02 ± 0.32 | 0.71 ± 0.40 | **-2.56 ± 0.39** |
| Slide (Å) | 0.18 ± 0.07 | -0.06 ± 0.13 | 0.74 ± 0.32 | **0.40 ± 0.08** | -0.05 ± 0.06 | -0.07 ± 0.14 |
| Twist (Deg) | **3.92 ± 0.45** | 0.02 ± 0.32 | **0.40 ± 0.08** | 6.87 ± 1.74 | 0.24 ± 0.43 | -1.50 ± 0.48 |
| Tilt (Deg) | 0.09 ± 0.21 | 0.71 ± 0.40 | -0.05 ± 0.06 | 0.24 ± 0.43 | **2.97 ± 0.55** | -0.82 ± 0.73 |
| Roll (Deg) | -0.76 ± 0.23 | **-2.56 ± 0.39** | -0.07 ± 0.14 | -1.50 ± 0.48 | -0.82 ± 0.73 | **3.90 ± 0.37** |
| **H1 Containing: Middle** | | | | | | |
| | Rise (Å) | Shift (Å) | Slide (Å) | Twist (Deg) | Tilt (Deg) | Roll (Deg) |
| Rise (Å) | **5.23 ± 0.03** | **-0.64 ± 0.12** | 0.11 ± 0.36 | **1.21 ± 0.24** | -0.74 ± 0.49 | 0.24 ± 0.08 |
| Shift (Å) | **-0.64 ± 0.12** | **2.24 ± 0.28** | 0.28 ± 0.21 | 0.05 ± 0.12 | -0.01 ± 0.06 | **-2.20 ± 0.30** |
| Slide (Å) | 0.11 ± 0.36 | 0.28 ± 0.21 | 1.25 ± 0.34 | **0.26 ± 0.05** | 0.11 ± 0.46 | -0.43 ± 0.12 |
| Twist (Deg) | **1.21 ± 0.24** | 0.05 ± 0.12 | **0.26 ± 0.05** | 1.45 ± 0.40 | -0.97 ± 0.42 | -0.14 ± 0.25 |
| Tilt (Deg) | -0.74 ± 0.49 | -0.01 ± 0.06 | 0.11 ± 0.46 | -0.97 ± 0.42 | **2.37 ± 0.25** | 0.07 ± 0.20 |
| Roll (Deg) | 0.24 ± 0.08 | **-2.20 ± 0.30** | -0.43 ± 0.12 | -0.14 ± 0.25 | 0.07 ± 0.20 | **2.91 ± 0.33** |
| **H1 Containing: Top** | | | | | | |
| | Rise (Å) | Shift (Å) | Slide (Å) | Twist (Deg) | Tilt (Deg) | Roll (Deg) |
| Rise (Å) | **2.05 ± 0.46** | 0.49 ± 0.17 | -0.15 ± 0.05 | **3.08 ± 0.69** | **-0.15 ± 0.04** | -0.68 ± 0.28 |
| Shift (Å) | 0.49 ± 0.17 | **3.82 ± 0.54** | -0.48 ± 0.13 | 0.39 ± 0.62 | 0.46 ± 0.16 | **-2.09 ± 0.36** |
| Slide (Å) | -0.15 ± 0.05 | -0.48 ± 0.13 | **0.45 ± 0.09** | -0.05 ± 0.07 | 0.08 ± 0.13 | 0.20 ± 0.10 |
| Twist (Deg) | **3.08 ± 0.69** | 0.39 ± 0.62 | -0.05 ± 0.07 | **6.64 ± 1.38** | -0.30 ± 0.09 | -0.70 ± 0.61 |
| Tilt (Deg) | **-0.15 ± 0.04** | 0.46 ± 0.16 | 0.08 ± 0.13 | -0.30 ± 0.09 | **1.07 ± 0.10** | -0.42 ± 0.16 |
| Roll (Deg) | -0.68 ± 0.28 | **-2.09 ± 0.36** | 0.20 ± 0.10 | -0.70 ± 0.61 | -0.42 ± 0.16 | **2.78 ± 0.40** |

Table S3: Kullback-Leibler divergence (KLD) values for two dimensional probability distributions of DNA motions between systems. Reference systems ($P(r)$ in eqn. 3) are given in the rows and the systems they are compared to ($P(r)$ in eqn. 3) are in the columns for each of three sets of distributions. For clarity, comparisons with low differences (KLD<3.3) are in blue, increased differences (3.3<KLD<6.7) are in green, high differences (6.7<KLD<10.0) are in orange, and very high differences (10.0<KLD) in red. The lower numerical values correspond to a greater similarity in probability distributions, whereas higher numerical values correspond to a greater dissimilarity.

| | α-Entry | | | | α-Exit | | | |
|---|---|---|---|---|---|---|---|---|
| | MonoNuc | MonoNuc with H1 | OctaNuc | OctaNuc with H1 | MonoNuc | MonoNuc with H1 | OctaNuc | OctaNuc with H1 |
| MonoNuc | 0.00 | 0.91 | 16.04 | 13.06 | 0.00 | 0.61 | 11.67 | 2.61 |
| MonoNuc with H1 | 0.41 | 0.00 | 13.63 | 10.69 | 0.56 | 0.00 | 12.76 | 3.67 |
| OctaNuc | 1.70 | 1.39 | 0.00 | 1.32 | 1.32 | 1.88 | 0.00 | 0.62 |
| OctaNuc with H1 | 1.46 | 0.96 | 3.46 | 0.00 | 0.45 | 1.25 | 2.48 | 0.00 |

| | β-Entry | | | | β-Exit | | | |
|---|---|---|---|---|---|---|---|---|
| | MonoNuc | MonoNuc with H1 | OctaNuc | OctaNuc with H1 | MonoNuc | MonoNuc with H1 | OctaNuc | OctaNuc with H1 |
| MonoNuc | 0.00 | 4.56 | 7.66 | 9.76 | 0.00 | 0.45 | 6.02 | 0.71 |
| MonoNuc with H1 | 6.94 | 0.00 | 13.07 | 11.76 | 0.33 | 0.00 | 3.26 | 0.09 |
| OctaNuc | 0.98 | 1.91 | 0.00 | 1.43 | 1.32 | 0.70 | 0.00 | 0.74 |
| OctaNuc with H1 | 1.60 | 1.19 | 2.02 | 0.00 | 0.48 | 0.09 | 3.09 | 0.00 |

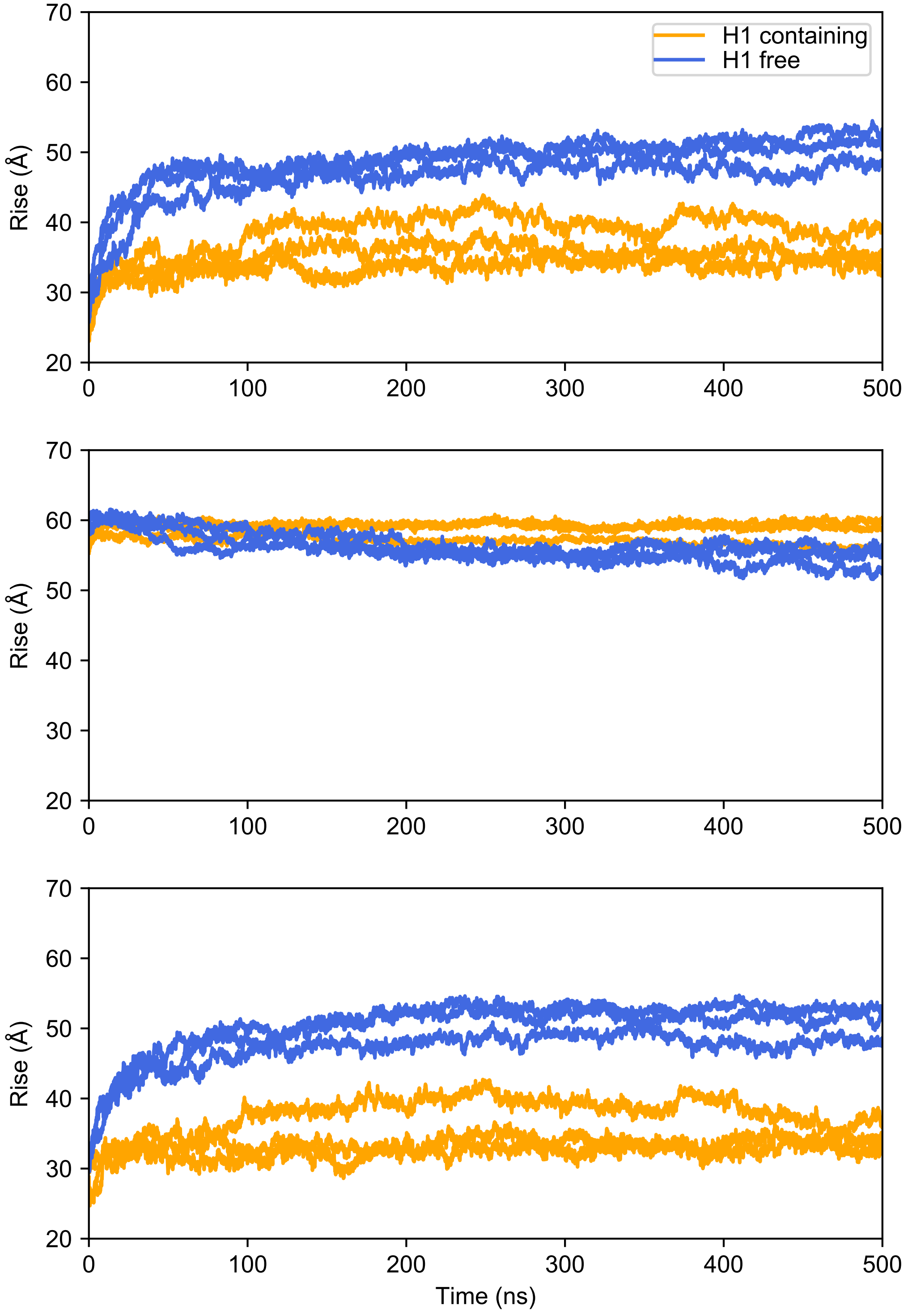


Figure S1: Nucleosomal rise for each simulation between the top, middle, and bottom four nucleosomes. Simulations containing H1 are shown in orange, simulations without H1 are in blue.

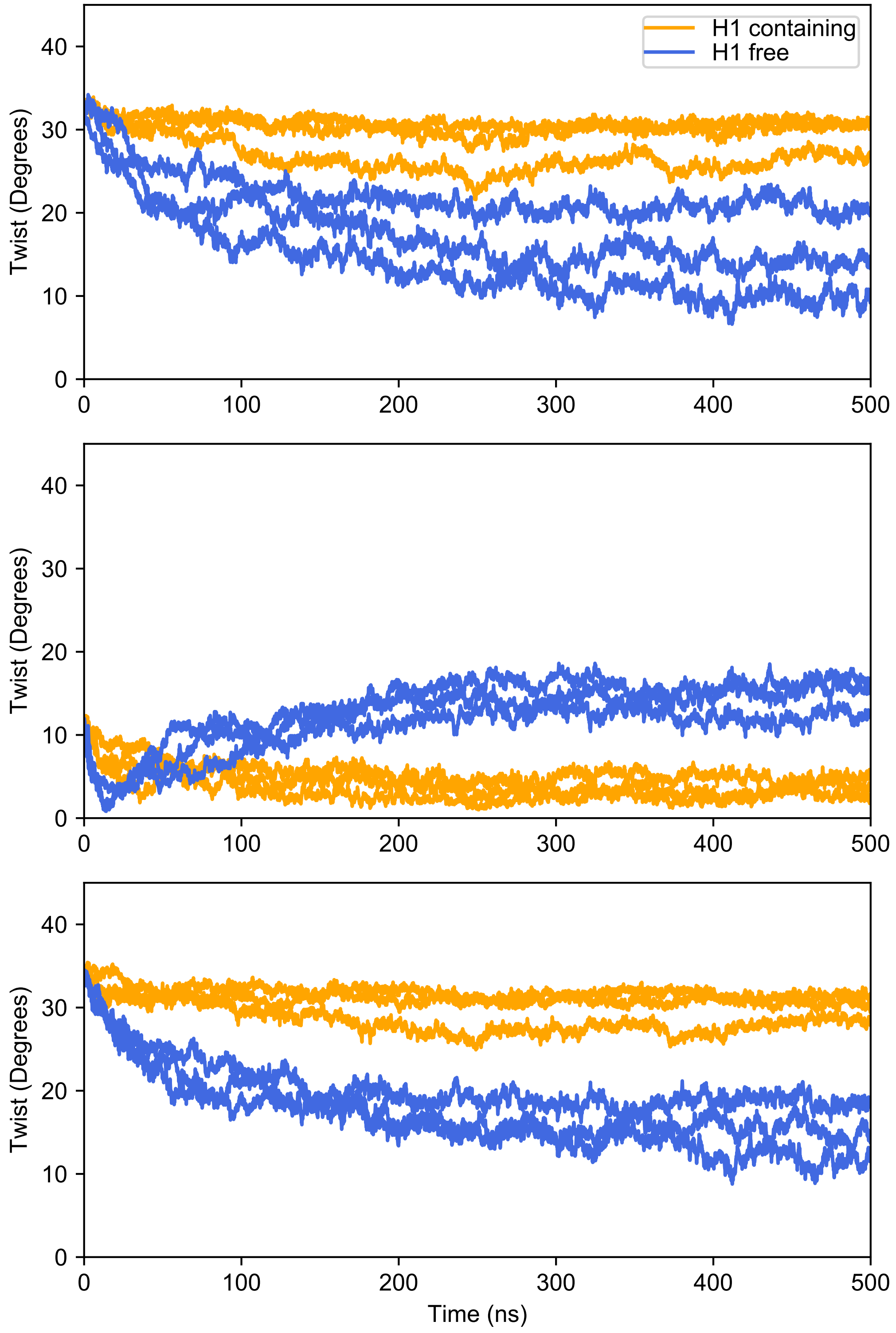


Figure S2: Nucleosomal twist for each simulation between the top, middle, and bottom four nucleosomes. Simulations containing H1 are shown in orange, simulations without H1 are in blue.

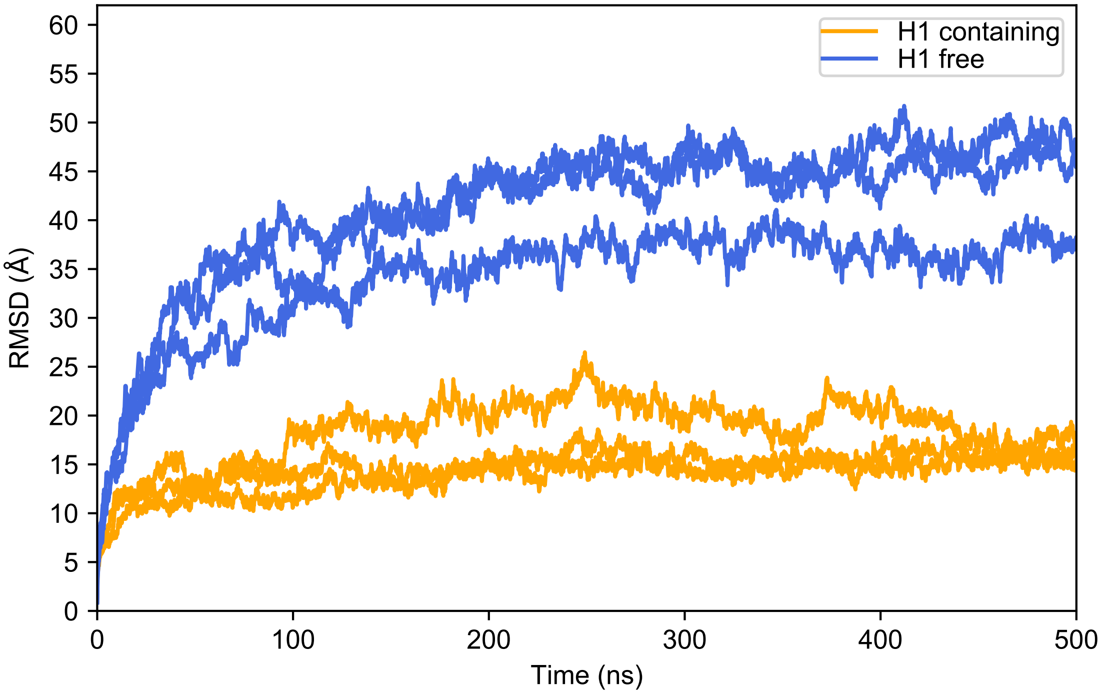


Figure S3: Root mean square deviations (RMSDs) for the protein $C_\alpha$ and DNA phosphate atoms for each simulation relative to the initial structure. Simulations containing H1 are shown in orange, simulations without H1 are in blue.

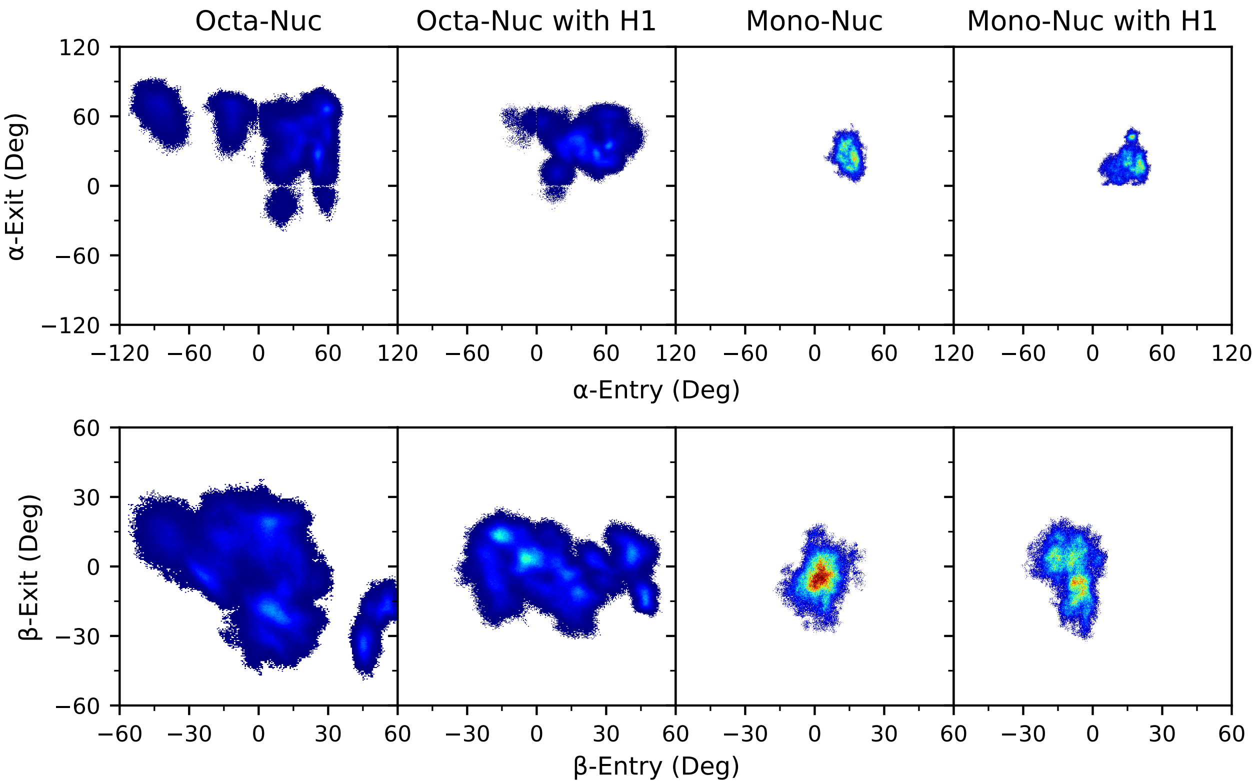


Figure S4: Entry vs. Exit sampling of linker DNA arms for alpha and beta angles. Density is represented as a gradient from blue (low density) to red (high density). Single nucleosome results are from previous work by Woods and Wereszczynski.[75]

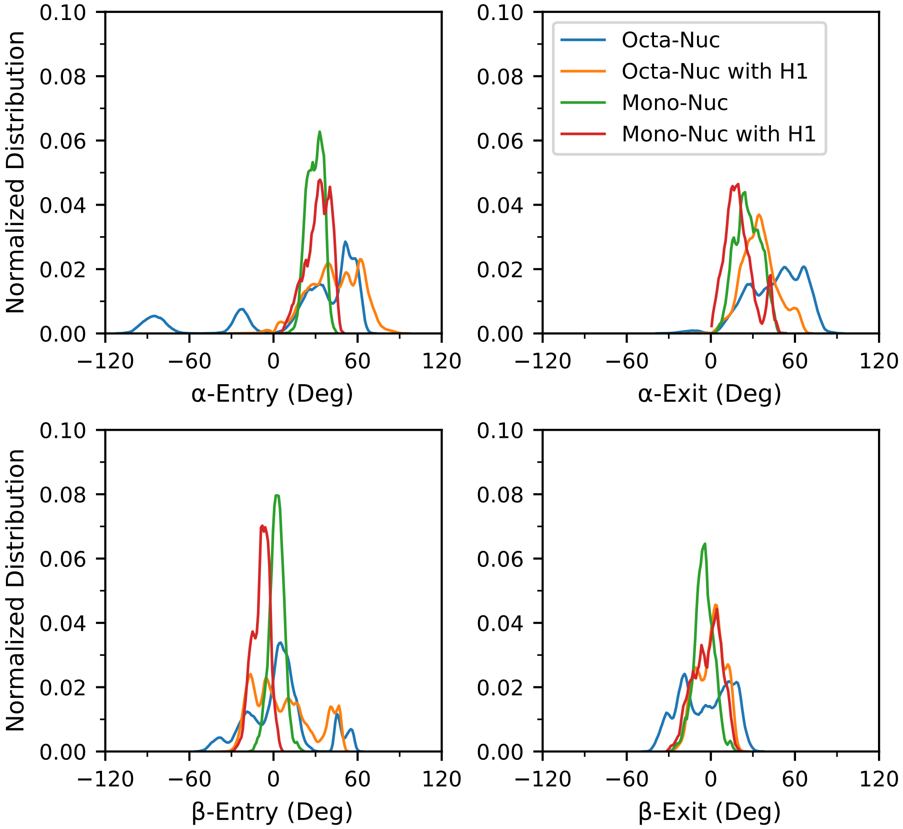


Figure S5: Alpha and Beta angle distributions for the octa-nucleosome with (blue) and without H1 (orange), and the mono-nucleosome with (green) and without H1 (red).

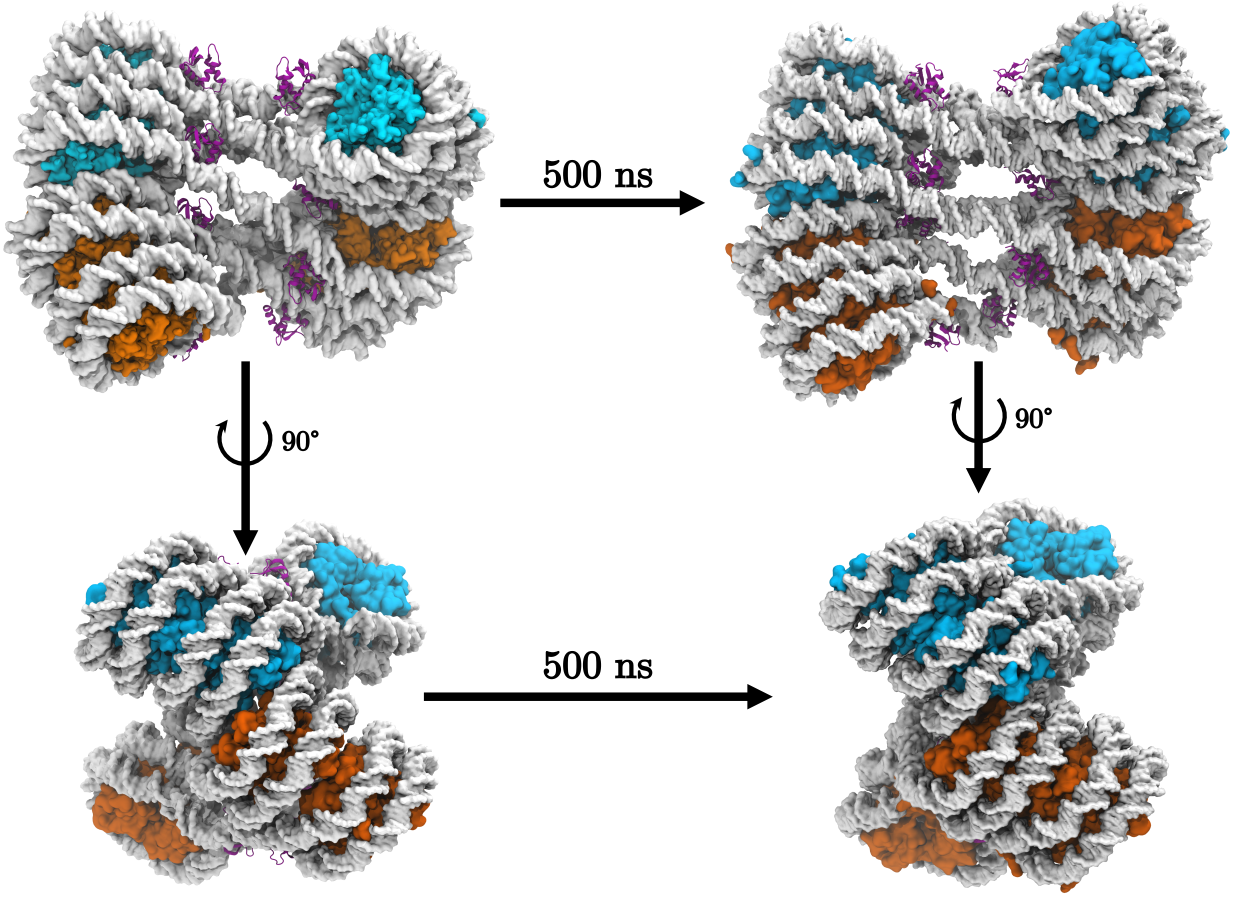


Figure S6: Shown are selected images from a system with the linker histone, at the beginning a simulation (left) and after 500 ns of production (right). The core histones in the poly-nucleosomes are colored to match Figures 1 and 3.

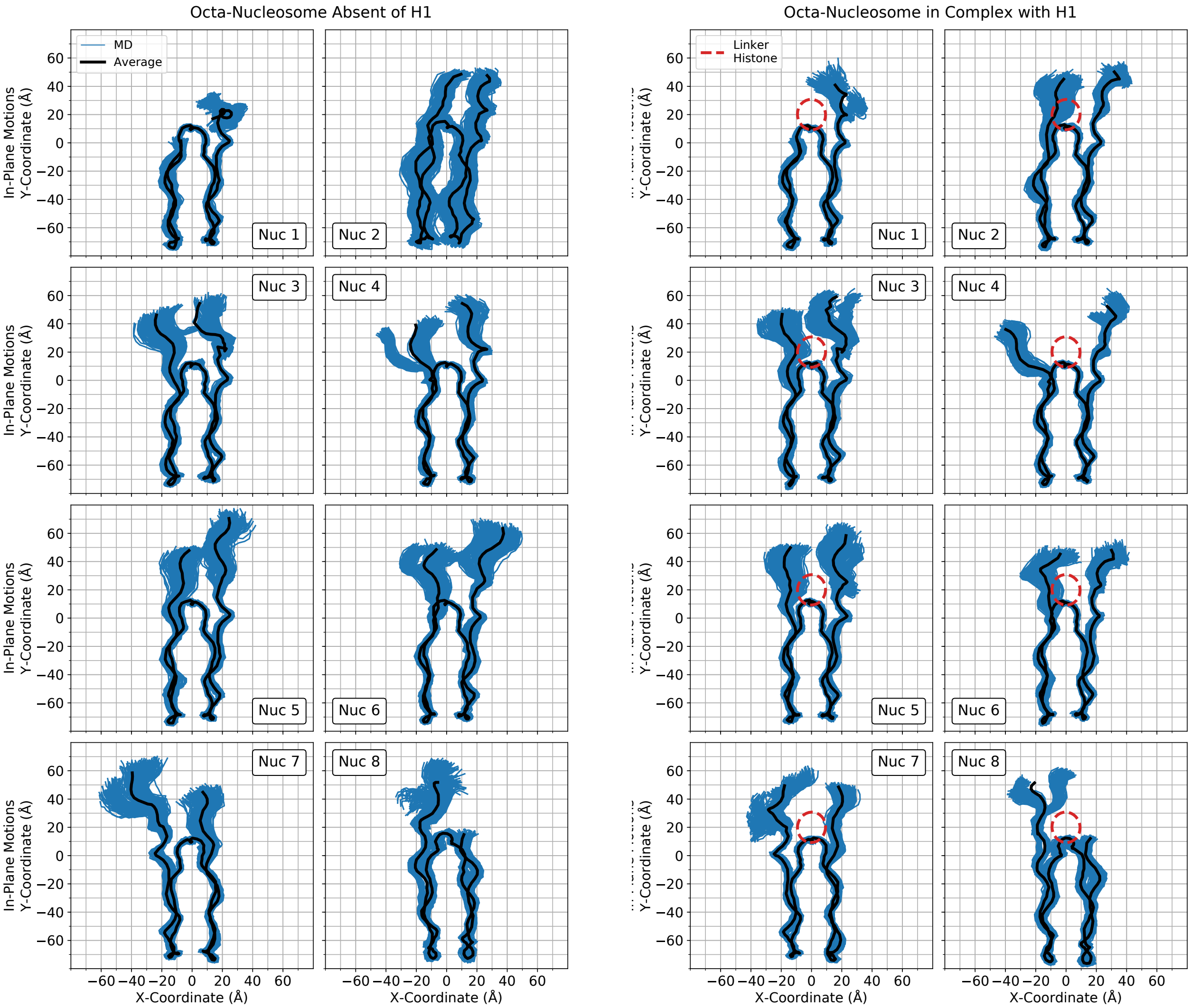


Figure S7: Out-of-plane (top) DNA motions sampled by the octa-nucleosome arrays absent (left) and in complex (right) with the linker histone H1. Shown in blue are configurations sampled throughout the MD simulation (263 representative frames - every 4 ns of simulation time) while the average configuration is shown in black. For reference, the relative position of each nucleosome in its array is labeled in the corner of each graph. This label is consistent with the numbering in Figure 3. Additionally, the approximate position of the linker histone is shown as a dashed-line red ellipse. Figures inspired by work from Shaytan *et al*[80] and single comparative nucleosome results were previously published by Woods and Wereszczynski.[75]

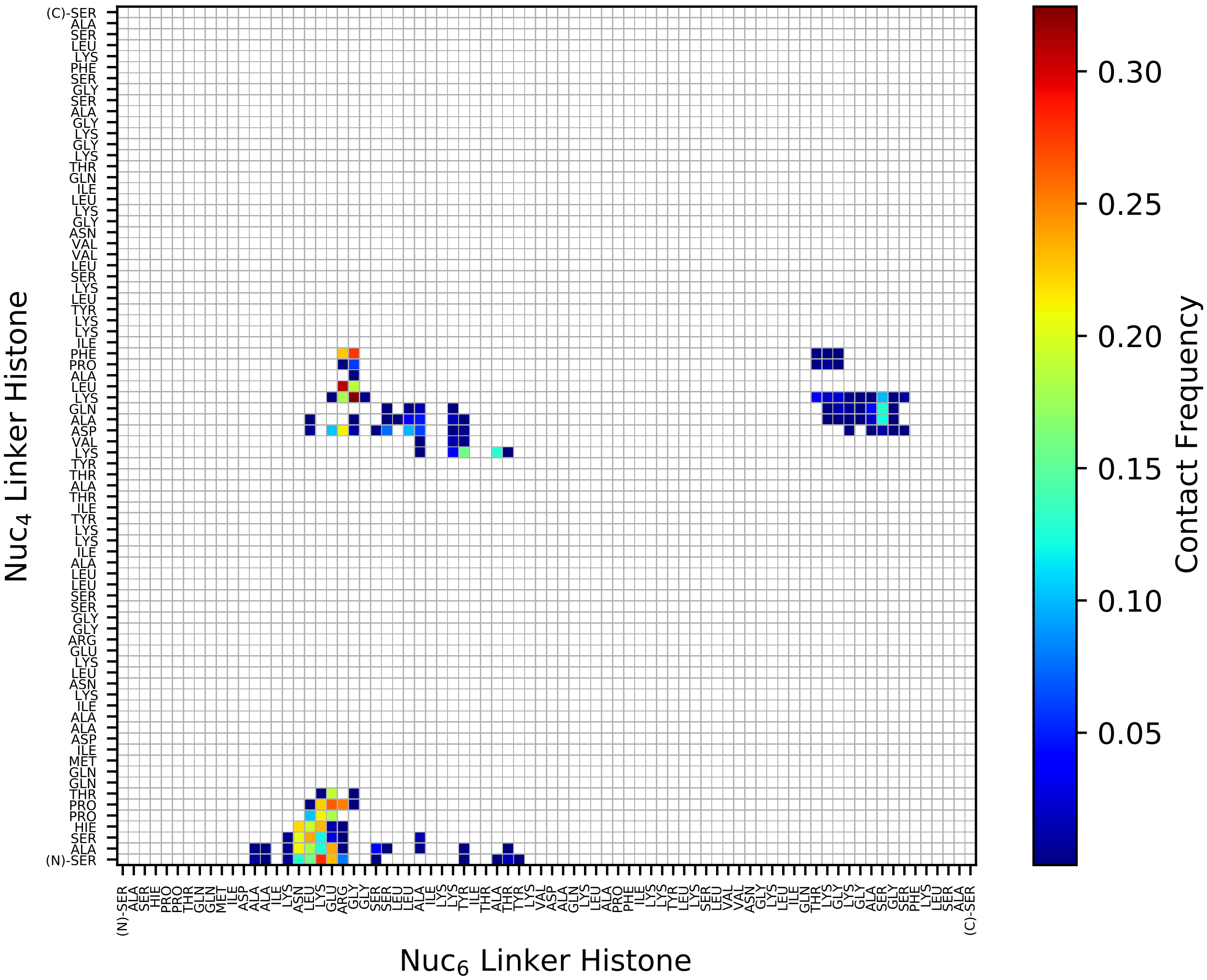


Figure S8: Contact frequency matrix between linker histones bound with $Nuc_4$ versus $Nuc_6$, respectively. Contacts were calculated using $C_\alpha$-atoms with a cutoff distance of 7.0 Å.

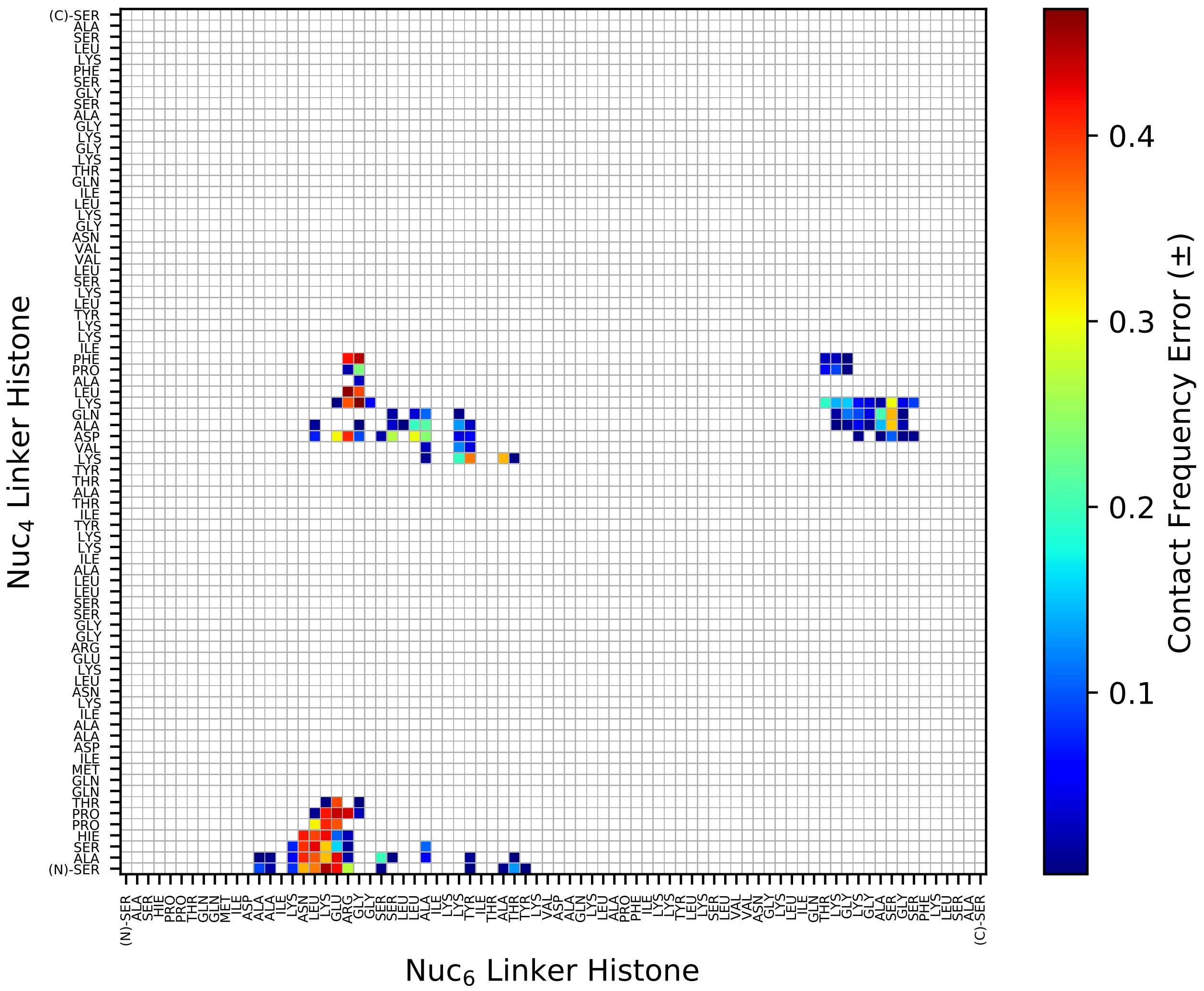


Figure S9: Contact frequency standard deviations matrix associated with Figure S8.

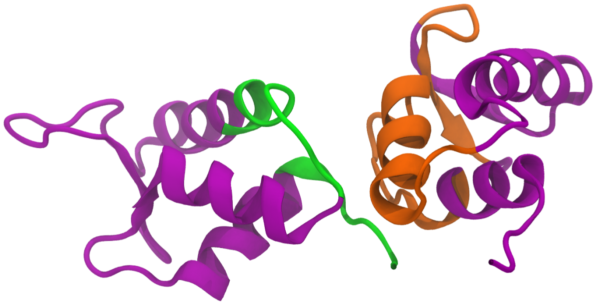

Figure S10: Contacting facets between linker histones associated with $Nuc_4$ (left) and $Nuc_6$ (right) as quantified in Figures S8 and S9. The colors, green and orange, represent regions of the proteins that were in contact at any duration throughout the simulations, respectively.
